# A Rapid Subcortical Amygdala Route for Faces Irrespective of Spatial Frequency and Emotion

**DOI:** 10.1101/078097

**Authors:** Jessica McFadyen, Martial Mermillod, Jason B. Mattingley, Veronika Halász, Marta I. Garrido

## Abstract

There is significant controversy over the anatomical existence and potential function of a direct subcortical visual pathway to the amygdala. It is thought that this pathway rapidly transmits low spatial frequency information to the amygdala independently of the cortex and yet this function has never been causally determined. In this study, neural activity was measured using magnetoencephalography (MEG) while participants discriminated the gender of neutral and fearful faces filtered for low or high spatial frequencies. Dynamic causal modelling (DCM) revealed that the most likely underlying neural network consisted of a subcortical pulvino-amygdala connection that was not modulated by spatial frequency or emotion and a cortico-amygdala connection that conveyed predominantly high spatial frequencies. Crucially, data-driven neural simulations demonstrated a clear temporal advantage of the subcortical route (70ms) over the cortical route (155ms) in influencing amygdala activity. Thus, our findings support the existence of a rapid functional subcortical pathway that is unselective of the spatial frequency or emotional content of faces.

## Introduction

The ability to rapidly detect external threats is essential to the survival of all species^1^. The amygdala has long been known to be involved in processing ambiguous and biologically-relevant stimuli^2^ but there is considerable controversy over how quickly the amygdala engages in visual processing^2^. One proposition born from converging human and animal evidence is that there is a short and direct colliculo-pulvinar pathway to the amygdala^3^. This so-called ‘low road’ is thought to transmit coarse visual information more rapidly than alternative ‘high roads’, which are thought to transmit fine-grained details via the visual cortex^4^. This multi-pathway proposition has sparked debate within the literature over the possible function and even the very existence of the low road^5–8^. The amygdala plays an important modulatory role in coordinating responses to biologically-relevant stimuli^9,10^ and in unconscious affective processing^3,11^. As such, it is essential that we understand the potential influence that a subcortical pathway might have over the earliest stages of numerous, cascading processes emanating from the amygdala.

Several studies have supported the anatomical existence of a subcortical pathway to the amygdala, as shown by diffusion imaging^12,13^, computational modelling^14,15^, and neuroanatomical tracing^16,17^. The estimated synaptic integration time of this relatively shorter pathway (estimated length: 58mm, estimated latency: 80–90ms) compared with the cortical visual stream (estimated length: 164mm, estimated latency: 145–170ms) suggests that the amygdala receives subcortical information faster than cortical information^18^.

This rapid subcortical pathway is thought to facilitate early processing of coarse visual information, such as low spatial frequency content, conveying the general “gist” of a visual scene to the amygdala^19^. Initial support for this notion came from a seminal functional magnetic resonance imaging (fMRI) study that found greater blood-oxygen-level-dependent (BOLD) signal in the superior colliculus, pulvinar, and amygdala to low spatial frequency fearful faces, while the extrastriate visual cortex showed greater BOLD signal for high spatial frequency faces^20^. These findings have recently been validated at the electrophysiological level, where fearful faces were found to evoke early activity (75ms post-stimulus onset) in the lateral amygdala but only when the faces were filtered for low spatial frequency^21^.

These studies do not address the critical element of causality, which is essential for investigating the influence of the subcortical route. While there is evidence that the neurons of the human superior colliculus are magnocellular^22^, little is known about the spatial frequency sensitivity of the human pulvinar^23^, although there is evidence that the pulvinar may be responsive to affective content^24,25^. Therefore, amygdala activation to low spatial frequency images should not be interpreted as being driven solely by a subcortical pathway. The fragility of this relationship is demonstrated by the often mixed findings as to whether low spatial frequency emotional stimuli are processed more rapidly^26^.

The aim of this study was to evaluate the causal direction of information flow along the cortical and subcortical pathways to the amygdala. Neural activity was measured with high temporal resolution using magnetoencephalography (MEG) while participants made gender judgements on faces filtered for different spatial frequencies. Dynamic causal modelling was then applied to these data to infer the direction of information transmission within hypothesised neural networks, using biophysically informed methods and Bayesian model comparison^27^. The hypothesised networks consisted of four increasingly complex models: 1) cortical connection from lateral geniculate nucleus to primary visual cortex (V1) to amygdala, 2) cortical connection plus a subcortical connection from pulvinar to amygdala, 3) cortical connection plus a medial connection from pulvinar to V1 to amygdala, and 4) cortical, subcortical, and medial connections. The medial connection was included to account for the possibility that the pulvinar influences amygdala activity indirectly via cortical sources^9^. After determining the most likely underlying neural architecture made up of a combination of these different processing streams, hypotheses were tested for the types of spatial frequency content propagated along each connection. Using this approach, we aimed to determine whether the subcortical pathway, if active, is modulated primarily by low spatial frequency images of biologically-relevant stimuli (i.e. neutral and fearful faces).

## Results

### Gender Discrimination Performance

Participants were instructed to report the gender of faces as quickly and accurately as possible. Faces were either unfiltered (i.e. broad spatial frequency) or filtered to contain only low or high spatial frequencies and either had a neutral or fearful expression. Outlier trials (i.e. trials where reaction time was more than 3 standard deviations above or below the mean for that participant) were removed from statistical analyses (*M* = 7.73%, *SD* = 4.24% of trials). Shapiro-Wilk tests revealed that the accuracy data was negatively skewed and thus Box-Cox transformations were conducted to satisfy conditions of normality. Hence, 2 (emotion) × 2 (spatial frequency) repeated-measures ANOVAs were conducted separately for reaction time (correct responses only) and the Box-Cox-transformed accuracy data. Greenhouse-Geisser values were used where appropriate and Bonferroni adjustments were made to correct for multiple comparisons. All reported statistics are significant at *p* < .05.

**Figure 1:**
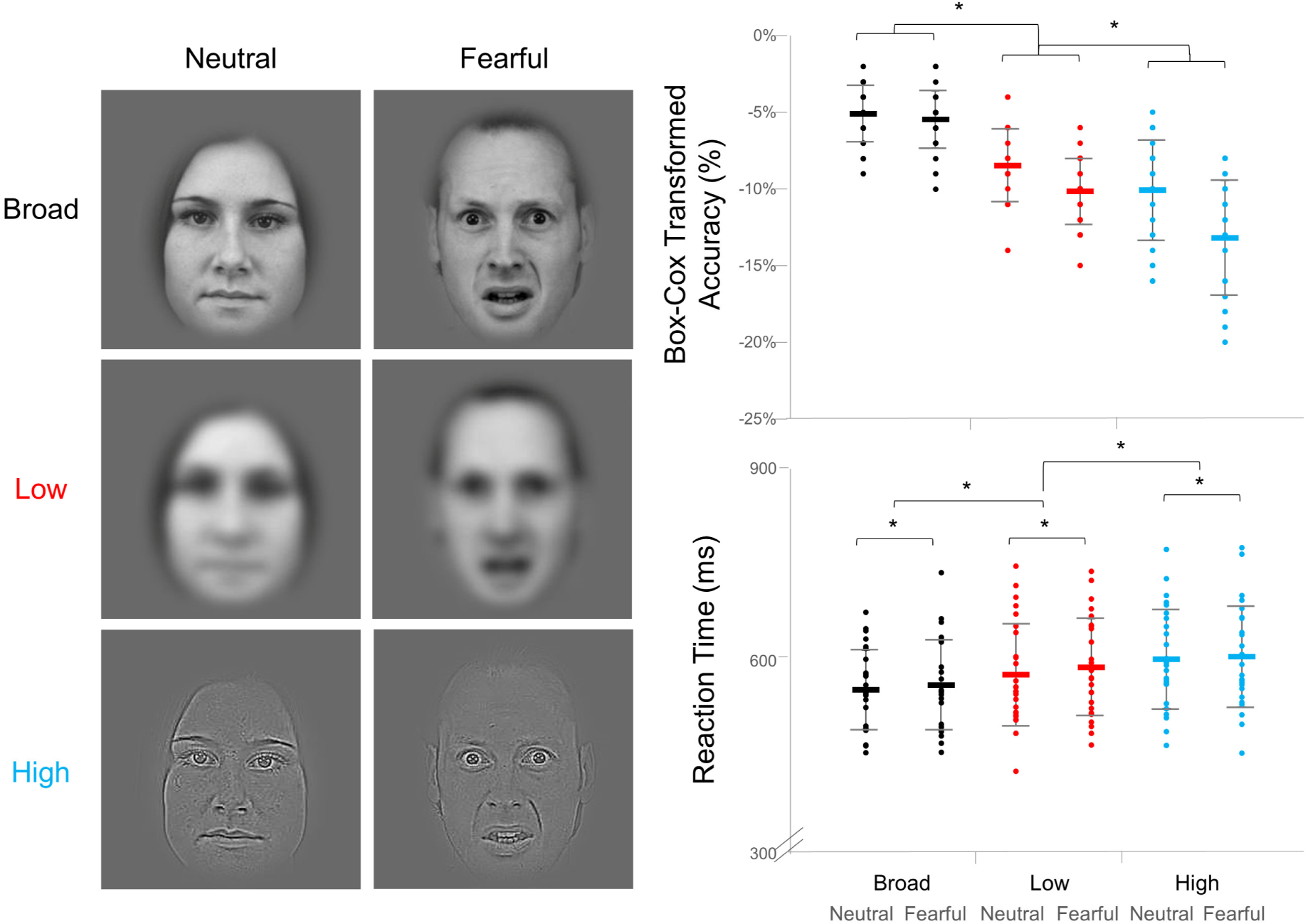
Experimental design and behavioural data. (*Left*) Examples of the face stimuli used in the experiment. The columns depict the Neutral and Fearful Emotion conditions and the rows depict the three spatial frequency conditions: Broad, Low, and High. (*Right*) Dot plots of each participant’s score for accuracy (top) and reaction time (bottom) in the gender judgement task. The accuracy data was Box-Cox transformed to satisfy conditions of normality. Black indicates Broad spatial frequency, red indicates Low, and blue indicates High. Each spatial frequency column contains a pair, where the left series represents Neutral expressions and the right series represents Fearful expressions. Standard error bars shown. * *p* < .05

Higher accuracy (*F*(1, 32) = 108.29, *p* = 6.65 × 10^−13^) and faster reaction time (*F*(2, 41) = 47.14, *p* = 2.30 × 10^−10^) were found for Broad spatial frequency faces (−5.30%, 559ms) than for Low (−9.29%, 585ms) spatial frequency faces and likewise for Low relative to High spatial frequency faces (−11.65%, 605ms). Significantly higher accuracy (*F*(1, 25) = 116.46, *p* = 6.74 × 10^−11^) and faster reaction time (*F*(1, 25) = 12.24, *p* = .002) were found overall for Neutral faces (−7.89%, 579ms), compared with Fearful faces (−9.60%, 587ms). An interaction was found in accuracy scores, such that the emotion effect was only present for Low and High spatial frequencies (*F*(2, 43) = 24.89, *p* = 2.33 × 10^−7^). Thus, gender discrimination was impaired more by the removal of low than high spatial frequencies. Furthermore, fearful faces slowed reaction times overall. Accuracy, however, was impaired by fearful expressions only when images were filtered for low and high spatial frequency.

Although gender discrimination performance has previously been thought of as equivalent between low and high spatial frequency faces^20,28,29^, our results support other studies that have found an advantage of low over high spatial frequency information^30–34^. Interestingly, few of these studies report an influence of emotional expression on performance^35^, which contrasts with our findings. The average luminance and root-mean-square contrast did not differ significantly between emotion conditions, as determined by separate 2 (emotion) × 3 (spatial frequency) repeated-measures ANOVAs, and so the difference was not likely due to low-level perceptual confounds. It is possible that fearful faces impaired performance because participants were instructed to respond as quickly (and as accurately) as possible, whereas such a time pressure may have been absent in previous studies.

### Spatiotemporal Analysis of MEG Sensor Data

Statistical parametric mapping was applied to neural data collected from 204 planar gradiometers. This sensor activity (field intensity) was converted into 3D maps of space (*x* and *y* mm^2^) and time (0–600ms, 5ms resolution). A 2 (emotion) × 3 (spatial frequency) within-subjects design was then modelled in a mass univariate analysis, correcting for multiple comparisons using Random Field Theory, as is typically performed in standard general linear model (GLM) analysis of fMRI data. The resultant statistical parametric maps were then compared between participants in a series of planned comparisons: 1) Low vs. High spatial frequency, 2) Neutral vs. Fearful Emotion, and 3) Interaction between Low and High spatial frequency and emotion. Broad spatial frequency was also used as an ecological control for Low and High spatial frequency in a set of two comparisons: Broad vs. Low and Broad vs. High (see **Supplementary Information** for a visualization of these results). All the following results for each comparison are *p* < .05, family-wise-error-corrected.

Significant differences in field intensity were found between Low and High spatial frequency across occipital, temporal, and central areas, spanning a time-window of 160–585ms (set-level *F*(2,25) = 38.86, *p* = 1.61 × 10^−6^; Fig. 2). Greater absolute field intensity for Low spatial frequency was found at the earliest significant time point (160ms) over right occipito-temporal areas and later on at 460 and 465ms over right temporal areas. All remaining clusters of significant activity showed greater field intensity for High than Low spatial frequency faces. These clusters were found at various time-points between 170 and 585ms, located bilaterally and centrally over occipital and parietal areas. Thus, we found distinct early (160ms) and late (460ms) components of Low spatial frequency processing despite overwhelmingly greater activity for High spatial frequency overall (170–585ms). Previous studies have also reported greater neural activity for Low than High spatial frequency images at approximately 160ms^31,34,36^. Moreover, similar findings have been reported for overall greater amplitude for High compared with Low spatial frequency faces^35,37,38^, with some studies reporting an earlier modulation than presented here (i.e. the M100 compared with the M170)^37,38^. This discrepancy may be due to differences in analysis technique, such that statistics on specific electrodes and time-windows are more specific (but also biased) compared with our more conservative and unbiased spatiotemporal analysis, where corrections for multiple comparisons are made across the entire sensor space and all time-points.

**Figure 2:**
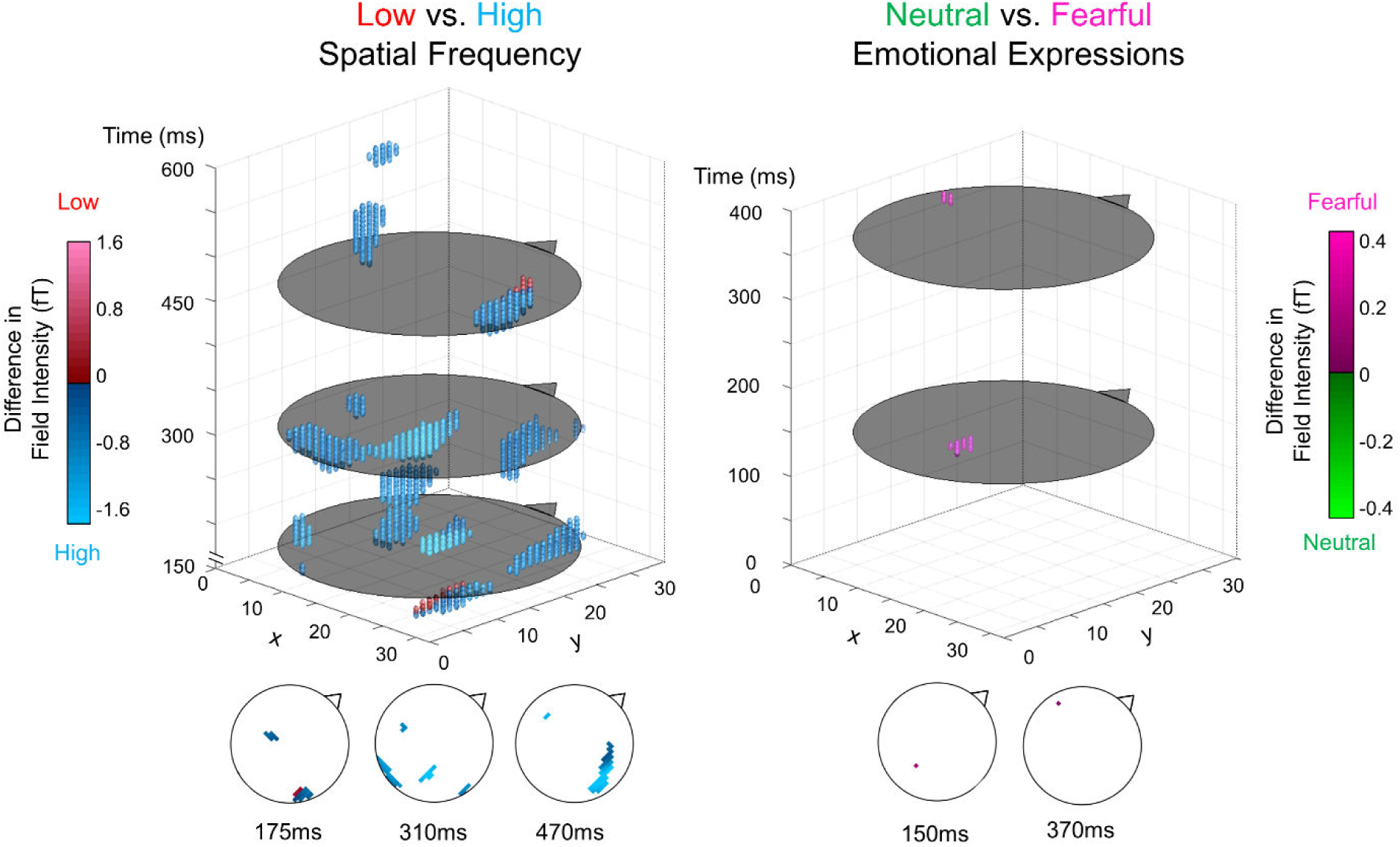
Statistical parametric maps of sensor data. A 3D representation of significant voxels across space and time. The flat grey circles in the 3D plot represent time-points of interest (for graphical purposes) which are each displayed as traditional scalp plots along the bottom of the figure (triangle/nose indicates faced direction). For Low vs. High spatial frequency (left), blue and red spheres indicate voxels that had significantly greater absolute field intensity for High and Low spatial frequency faces, respectively. Similarly, for Neutral vs. Fearful on the right, pink and green indicate greater absolute field intensity for Fearful and Neutral expressions, respectively. All points are significant at *p* < .05, family-wise error corrected.

Activity elicited by Fearful faces was significantly greater than by Neutral faces at two distinct clusters: an occipital peak at 150ms and a left temporal peak at 370ms (set-level *F*(1,25) = 38.48, *p* = 1.73 × 10^−6^), indicating typical early and late emotion effects as reported in the literature^39^. This effect did not, however, interact significantly with spatial frequency. Thus, although Low spatial frequency processing was indeed found to be faster overall, this did not result in enhanced processing of fearful faces. Hence, our findings do not support an automatic prioritisation of Low spatial frequency fear processing, which is in line with the findings of a recent meta-analysis^26^.

### Assessing Neural Latency with Cross-Correlation

In the spatiotemporal analysis, the earliest significant difference between spatial frequency activation was found at 160ms, where field intensity was greater for Low than High spatial frequency, followed by significantly greater activity for High than Low at 170ms onwards. To determine whether this apparent temporal difference was significant, cross-correlation analyses were performed. This approach assumes that two paired waveforms (i.e. Low and High spatial frequency ERFs at each sensor) are highly correlated, which was indeed the case in this dataset (average *R*^2^ across channels and subjects = 82.79%). Thus, we computed the relative lag between these waveforms at each of the 204 sensors. This resulted in 96 (47.06%) of sensors with a significant lag (Bonferroni-corrected, *p* < 2.45 × 10^−4^) between Low and High spatial frequency faces, where all 204 sensors demonstrated an earlier effect of Low spatial frequency (*M* = 7.13ms, *SD* = 2.19ms; Fig. 3). This finding is supported by similar studies showing earlier M170 latencies for Low compared with High spatial frequency image processing^34,35^. Therefore, there appears to be a significant temporal disadvantage for High compared with Low spatial frequency faces, such that the waveform as a whole (which encompasses multiple face processing components like the M100 and M170) is shifted later in time.

**Figure 3:**
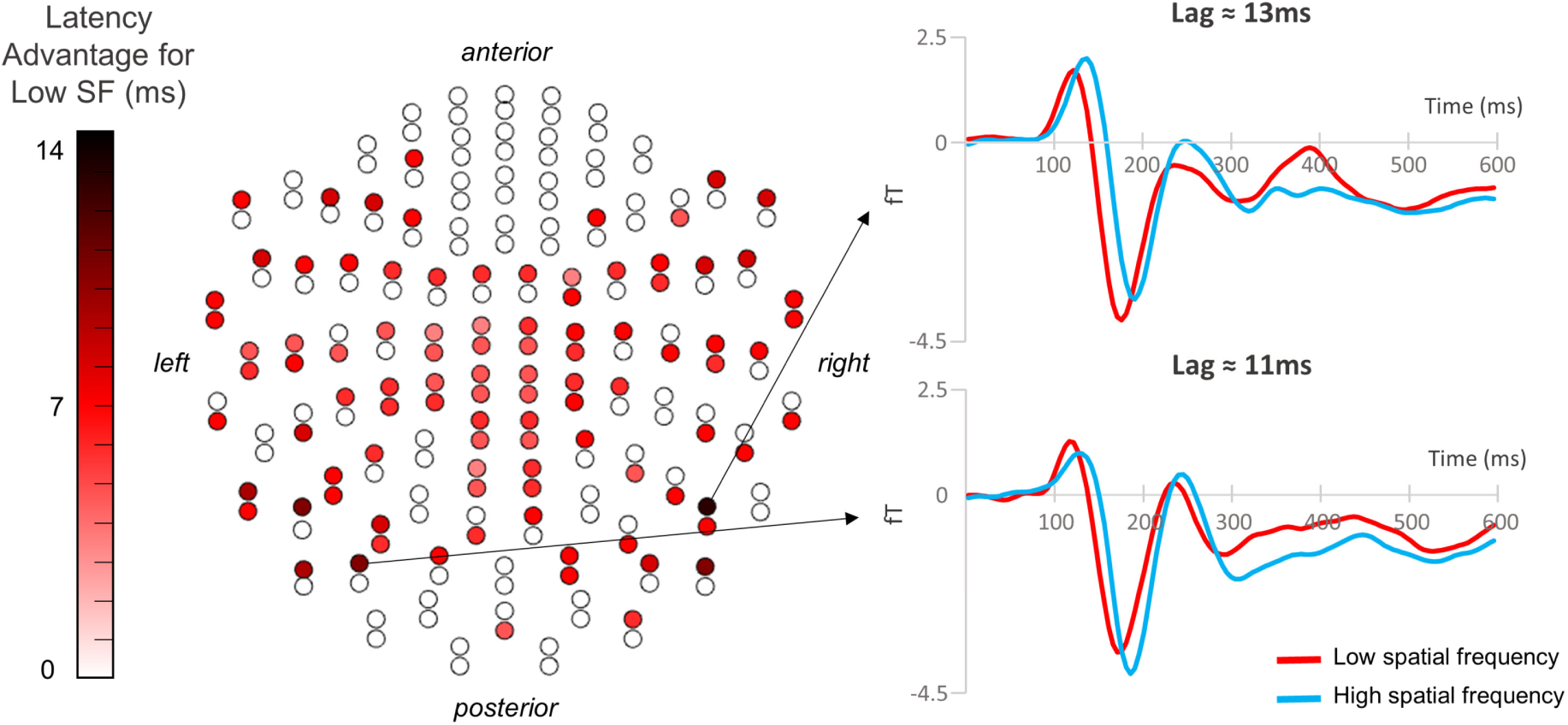
Latency advantage for low spatial frequency faces. (*Left*) A sensor map displaying 204 planar gradiometers, where red indicates a sensor that showed a significant time difference between Low and High spatial frequency, such that darker red indicates an earlier neural effect of Low compared with High spatial frequency faces. (*Right*) Activity at two example sensors that showed the greatest time difference.

### Dynamic Causal Modelling of Neural Networks

Dynamic causal modelling is a biophysically informed method of comparing the likelihood of hypothetical neural networks that underlie a given dataset, compromising between model accuracy and complexity^27^. This technique was used in two stages: an *anatomical* stage, where we compared the likelihood that cortical, medial, and subcortical connections were recruited across all conditions, and a *functional* stage, where we took the winning model from the previous stage and compared the likelihood that connections were modulated by spatial frequency and/or emotion.

Sources were modelled using equivalent current dipoles based on validated functional coordinates from previous literature (MNI coordinates: LGN – left [−22, −22, −6], right [22 −22 −6]; V1 – left [−7 −85 −7], right [7 −85 −7]; pulvinar – left [−12 −25 7], right [12 −25 7]; amygdala – left [−23 −5 −22], right [23 −5 −22])^14^. Input was modelled as entering the LGN and pulvinar and only the first 300ms of activity post-stimulus onset was modelled so as to capture early visual processing, specifically.

In the anatomical stage, four families of models were constructed (Fig. 4). Each family consisted of models for each possible combination of forward and backward connections, resulting in four models for the Cortical family, eight models for Dual, eight models for Medial, and sixteen models for All. Each model contained nodes for the left and right hemispheres but cross-hemisphere connections (e.g. from the left amygdala to the right amygdala) were not modelled.

**Figure 4:**
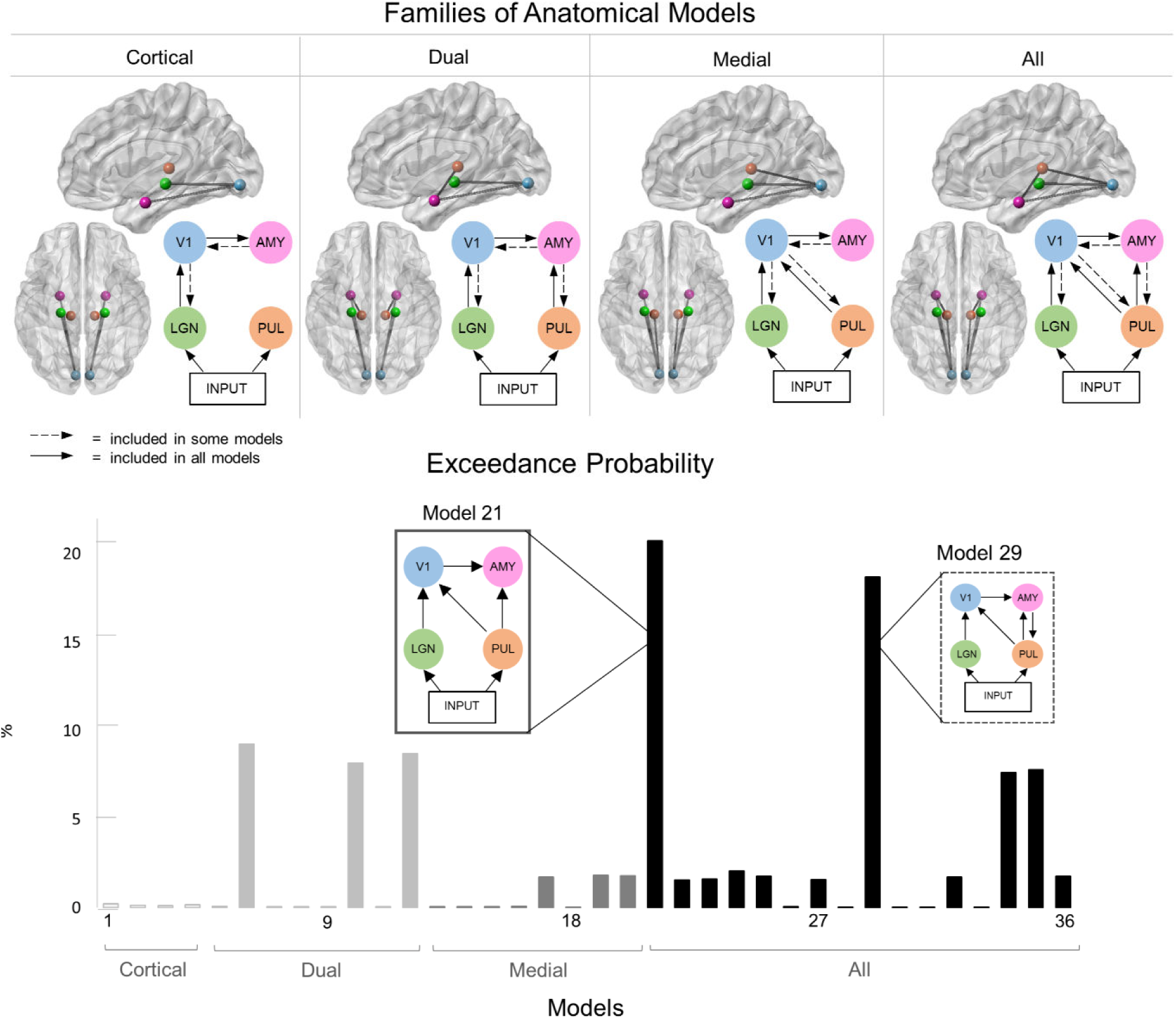
Selection of families of anatomical models. (Top) Four families of anatomical models were tested. Green nodes represent the LGN, blue the primary visual cortex (V1), pink the amygdala (AMY), and orange the pulvinar (PUL). The 3D brain images (saggital and axial views) are shown. Note that all models consisted of separate left and right hemispheres. (Bottom) The exceedance probability of each model, grouped into the four families (light grey for Cortical, grey for Subcortical, dark grey for Medial, and black for All). Model 21, which had the highest probability, is shown in the left break-out box, while the next most probable model, Model 29, is shown in the right break-out box.

Random effects Bayesian Model Selection was conducted on all models grouped into families. The greatest exceedance probability was found for the All family (59.61%) compared with the Dual (23.69%), Medial (13.29%), and Cortical (3.40%) families. Therefore, it is clear that the inclusion of the subcortical route (i.e. the All and Dual families) vastly increased model likelihood. Within the winning All family, two models had comparably higher exceedance probability than the other fourteen: Model 21 (20.72%) and Model 29 (18.69%). Model 21 was the simplest in the family of models, containing only forward connections, whereas Model 29 was only incrementally more complex due to the addition of a backward connection between the amygdala and pulvinar. Since previous research has established that backward connections contribute more to a model’s explanatory power as peristimulus time increases^40^, we speculate that the below-chance difference between these two models (chance being 100% divided by the number of models in the space, giving 2.78%) may be because the backward connection’s effect only occurred in the latter part of the 0–300ms time-window. Applying Occam’s razor, we selected Model 21 as the most parsimonious explanation for our data because this model had the fewest connections and also the greatest exceedance probability. Hence, the anatomical foundation for the following series of functional models consisted of the well-known cortical stream, the controversial subcortical pathway, and the hypothesised medial connection between the pulvinar and the visual cortex.

The functional stage was designed to determine the likelihood of different routes of modulation by spatial frequency, emotion, or both, using a between-trial effects approach (i.e. differences between Low and High spatial frequency or between Neutral and Fearful emotional expressions)^41^. First, we determined the possible paths along which visual information may be modulated (four possibilities: V1-AMY, LGN-V1-AMY, PUL-AMY, PUL-V1-AMY). These modulatory pathways were then combined in every possible way, resulting in nine models each for modulation by spatial frequency, emotion, or both (Fig. 5). Thus, the entire model space consisted of 28 models, including a null model precluding any modulation.

**Figure 5:**
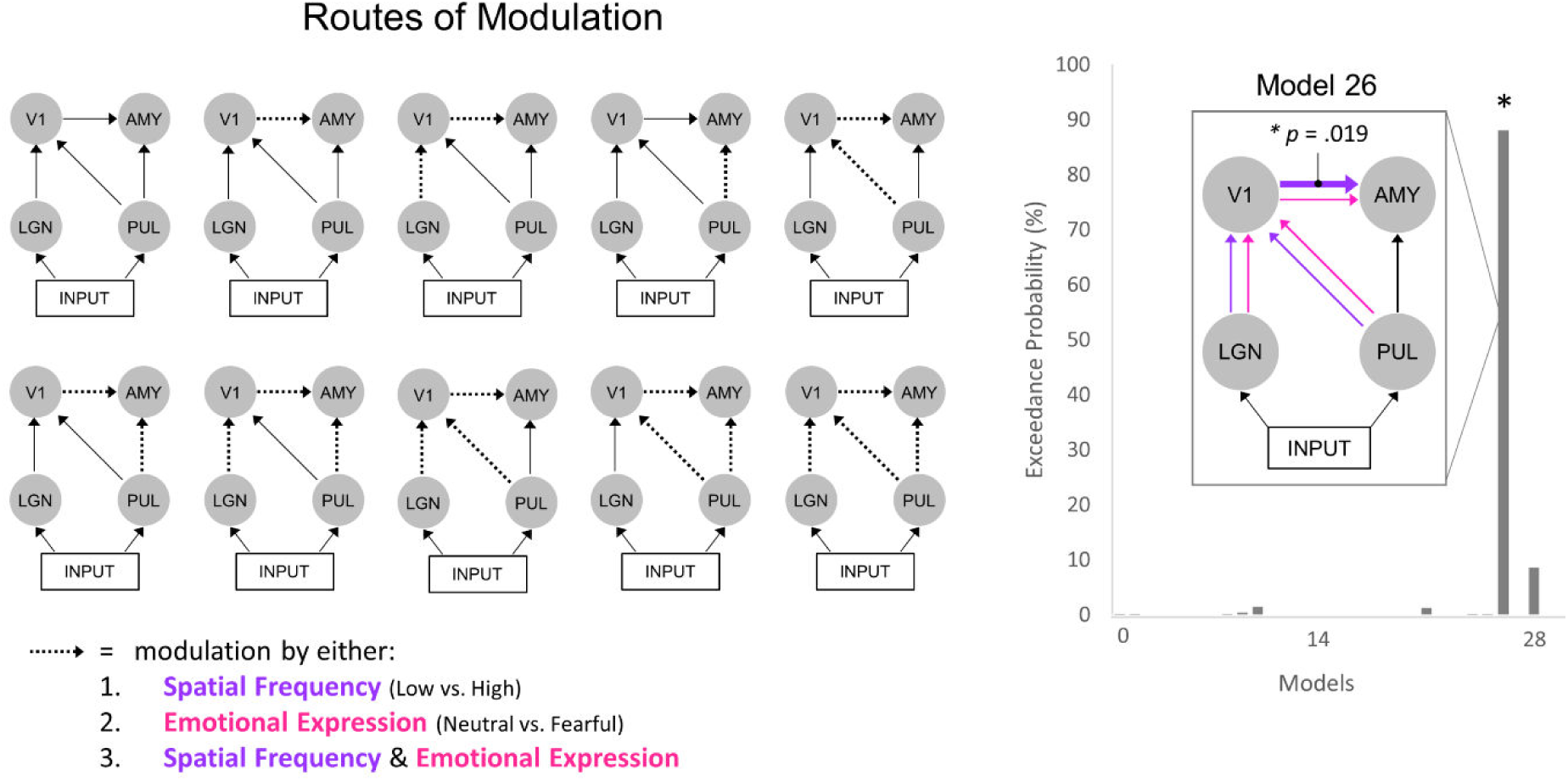
Routes of modulation by emotion and spatial frequency. (Left) Each diagram depicts a plausible route of modulation, where the dotted arrows indicate potential modulation by spatial frequency, emotion, or both. Note that the first model in the upper left is the Null model with no modulated connections. The other nine models were each uniquely modulated. (Right) The exceedance probability, determined via Bayesian Model Selection, for all 29 models. The model shown in the break-out box is the winning model (Model 26), where all connections were modulated by spatial frequency and emotion except the subcortical route. Classical t-tests revealed that only the parameter estimates for the cortical connection were significantly greater than zero (*p* = .019).

Random effects Bayesian Model Selection revealed that Model 26 had the greatest exceedance probability (88.04%), demonstrating that all pathways except the subcortical pathway were likely modulated by both spatial frequency and emotion within the given dataset. To infer the generalizability of this neural network to the population, classical one-sample t tests were conducted on individual parameter (i.e. connection) estimates. After Bonferroni correction for multiple comparisons, only the modulation of the cortical connection from primary visual cortex to amygdala was significantly greater than zero (*p* = .019).

### Sensitivity Analysis

Having found a clear winning model for both the underlying neural architecture and the likely modulation of parameters by spatial frequency and emotion, we investigated the direction of information flow in the network (i.e. was V1-amygdala activity greater for high or low spatial frequency?) and also the timing of flow within the network (i.e. did the subcortical pathway perturb amygdala activity earlier than the cortical pathway?) To do this, we employed sensitivity analysis, which uses simulated data generated by an estimated dynamic causal model inferred from the data. This analysis technique tests the influence of particular parameters within a network by artificially amplifying coupling strength through stimulation. If the parameter influences the activity at a particular node in the simulated network, then it should be possible to observe a fluctuation in the simulated activity^42–44^. Therefore, using a data-driven modelling approach, we were able to make inferences about the influence of the subcortical, cortical, and medial connections on visual cortex and amygdala source activity.

A replica of the wining model was constructed to split the modulatory effects of spatial frequency and emotion at the cortical and medial streams into Low-Neutral, Low-Fearful, High-Neutral, and High-Fearful (each indicating spatial frequency and emotional expression, respectively), allowing us to observe interaction effects. Analyses were conducted separately on the anatomical presence and functional modulation of each connection. Waveforms of amygdala and V1 source activity were produced for perturbation by each connection type. The entirety of each of these waveforms was statistically analysed by creating statistical parametric maps (*source activity* × *time*) and using Random Field Theory to correct for multiple comparisons across time (*p* < .05), similar to the previous spatiotemporal analysis.

First, we investigated how each anatomical connection influenced V1 and amygdala activity by boosting the coupling parameters for each connection. V1 source activity was equally influenced by its forward connections from lateral geniculate nucleus and pulvinar from 110–160ms (Fig. 6, **left**). Changes in amygdala activity, on the other hand, were significantly greater for the subcortical pathway than for all other connections between 70–90ms and 110–165ms (Fig 6., **centre**). There was also a significantly greater change in amygdala source activity for the direct cortical connection from V1 than from the more indirect LGN-V1 (155–190ms) and pulvinar-V1 (165–180ms) connections.

**Figure 6:**
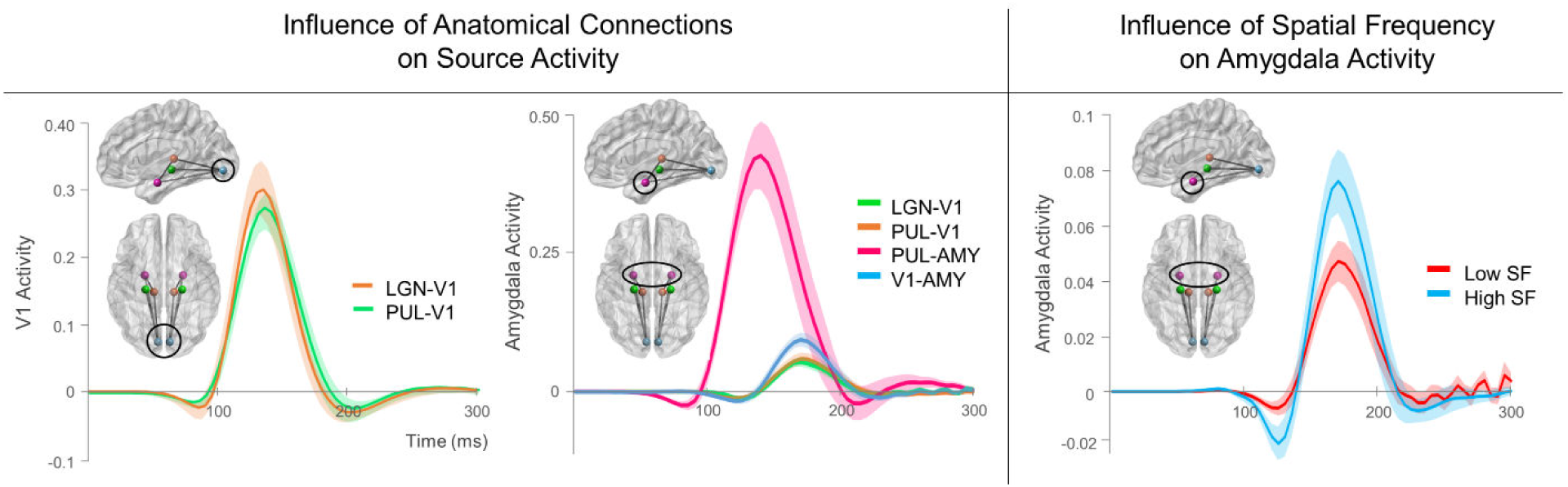
Sensitivity analysis of connection type on source activity. Sensitivity Analysis of anatomical connectivity (left, centre) and modulatory (right) connections. Each wave depicts the simulated perturbation in source activity by either the presence of a connection, as in the left-hand and centre graphs, or the modulation by Low or High spatial frequency, as in the right-hand graph.

To determine whether these apparent time differences in amygdala perturbation were significant, cross-correlation was conducted on pairs of the four connections. As expected, the perturbation by the subcortical pathway at 70ms was significantly earlier than the perturbation by the LGN-V1 (by 33.27ms, p = 6.638 × 10–10), PUL-V1 (by 31.54ms, p = 1.956 × 10–12), and V1-AMY connections (by 32.88ms, p = 9.659 × 10–13), each of which perturbed activity later at approximately 155ms.

A separate sensitivity analysis was conducted on the functional modulation of connections in the winning model. This first revealed that neither spatial frequency nor emotion modulation influenced V1 source activity through LGN or pulvinar input pathways. Interestingly, a main effect of Spatial Frequency was found for amygdala source activity, showing greater perturbation by High than Low spatial frequency during 125–130ms and 165–180ms (Fig. 6, **right**). Upon observation of each individual connection (i.e. LGN-V1, V1-AMY, and PUL-V1), it appears that High spatial frequency yielded a greater general effect across connections but that this effect was predominantly driven by modulation along the cortical V1-AMY connection, which was the only significant simple effect. This corroborates the significant *t* statistic on this parameter found within the winning functional Model 26.

## Discussion

We have presented computational evidence supporting the existence of a subcortical route to the amygdala that facilitates rapid visual information transfer as early as 70ms, irrespective of spatial frequency or emotional content. Participants viewed low, high, and broad spatial frequency faces exhibiting neutral or fearful expressions. Performance on a gender discrimination task was better for low than high spatial frequency faces and for neutral than fearful faces, while neural activity was greater and slower overall for high compared with low and broad spatial frequency faces. Taken together, this suggests that high spatial frequency faces were more computationally demanding^45^.

Dynamic causal modelling was used to test hypotheses for the most likely structure and function of the neural network underlying face perception, particularly the rapid processing that occurs within the first 300ms. The anatomical stage demonstrated that the most likely network consisted of forward-only cortical, subcortical, and medial connections to the amygdala, providing evidence for the existence of the ‘low road’ to the amygdala that operates in parallel with the ‘high road’, as shown in previous dynamic causal modelling studies^14,15^. In the winning model from the functional stage, the cortical and medial pathways were modulated by spatial frequency and emotion but the subcortical pathway was not, indicating that activity along this pathway was indistinguishable between conditions. Classical statistics on the parameters themselves revealed that only the spatial frequency modulation of the cortical connection was consistent across participants and thus generalizable to the population. Sensitivity analysis on a more detailed replica of the winning model demonstrated that there was significantly earlier perturbation of amygdala activity by the subcortical route (70ms) than by the cortical or medial pathways (155ms), regardless of emotion or spatial frequency content.

### Multiple Parallel Pathways

Our results lend support to anatomical evidence of this pathway^12,17,46^, as well as previous demonstrations of a functional subcortical route to the amygdala using dynamic causal modelling of MEG data^14,15^. This clearly illustrates the important causal role of the pulvinar in triggering amygdala activity. Importantly, we also found evidence for a functional medial connection from the pulvinar to the visual cortex, previously demonstrated in human and non-human primates^47^. Such a cortico-pulvinar connection has been suggested to mediate the subcortical pathway, thus rendering it *indirect*^6^. Our results dispute this alternative explanation, as we found the medial connection to be forward-only and its influence on amygdala activity was estimated to be over 30ms later than the subcortical route’s influence. Our results therefore support a multiple, parallel processing account of visual information transfer to the amygdala, within which there exists a rapid subcortical pathway^11^. This adds significant weight to the prospect that the subcortical route plays a role in unconscious, or pre-attentive, affective processing by providing a rapid direct pathway to the amygdala that operates independently of the cortex^11,48^.

### The Subcortical Route is “Blind” to Spatial Frequency

Contrary to our initial hypothesis, the subcortical pathway was modulated neither by spatial frequency nor emotion. This lack of modulation of the subcortical route by spatial frequency was somewhat unexpected given previous fMRI findings for increased low spatial frequency activity in the superior colliculus, pulvinar, and amygdala during emotional face viewing^20^, as well as the widely-held assumption that these subcortical areas consist of magnocellular neurons^49^. Furthermore, a recent intracranial EEG study by Mendez-Bertolo et al. found lateral amygdala responses to fearful faces that were specific to low spatial frequencies^21^.

One potential explanation for these discrepancies is that our stimuli were matched for both luminance and contrast, unlike other studies where only luminance was matched^20,21^. The effects of contrast equalisation on spatial frequency processing are particularly important during early (i.e. < 100ms) visual processing^35^. Supporting this notion is another intracranial EEG study that used luminance-and contrast-equalized stimuli but did not observe significant differences between low and high spatial frequency visible faces until 240ms^50^. Thus, teasing apart the effects of contrast and spatial frequency on subcortical activity may be a promising avenue for future research.

Another potential explanation for these apparently conflicting results is that they simply view neural processing at different scales. A multitude of evidence supports accurate localisation of deep sources in MEG^51,52^, especially when source reconstruction techniques are used in conjunction with coordinate locations known *a priori* from fMRI studies employing the same paradigms^14,51,53^ such as in the present study. Combining this with a dynamic causal modelling approach allowed us to consider the likelihood of the broad underlying neural architecture at the *systems* level and to make inferences on the causal nature of information flow along neural connections. This critical element of causality was absent in previous fMRI research, leaving open the possibility that the spatial frequency effects they reported were driven by feedback connections from other neural areas^20^.

Electrophysiological approaches, on the other hand, yield highly spatially and temporally specific effects at the *cellular* level^50,54^. This technique has revealed early spatial-frequency-and emotion-specific effects at the level of single cell populations within the amygdala^21^. The present findings, however, extend this result by demonstrating that these cellular differences may be less meaningful when examined at the neural systems level, where we found a higher likelihood of an *unmodulated* subcortical route than one modulated by spatial frequency or emotion. The complementary nature of these findings is further supported by the similar latencies found for amygdala perturbation (approximately 70ms). Hence, while our study provides causal evidence to support the claim that this early perturbation in the lateral amygdala was driven by the subcortical route^21^, small short-lived differences in spatial frequency processing may not have been sufficient to be detected (and thus may not be meaningful) at a broader systems level.

### Rapid Transmission of both Neutral and Fearful Faces

Emotional expression modulated activity along the medial pulvinar-V1 pathway but not along the subcortical pathway. This result is consistent with previous dynamic causal modelling studies that also report a lack of emotional modulation along the subcortical pathway^14,53^. Indeed, we would expect that significant emotional responses to fearful faces would not be observed until visual information is processed by the amygdala or other higher-order neural areas^15,55^. This is supported by the latency of the emotion effects we observed at the sensor level (occurring at approximately 150ms). Some degree of pre-emptive processing in the pulvinar, however, is supported by previous research on pulvinar response-specificity to faces^24,56^ and snakes^24,57,58^. There has been debate, though, over whether these pulvinar responses are generated independently or are a result of cortical mediation^6,8^. Our results suggest that the pulvinar is independently capable of some degree of filtering of facial expressions, as shown by the modulation within the winning model.

### A Generalised Functional Role of the Subcortical Route

Despite the capability of the pulvinar in filtering both spatial frequency and emotional content, we found the subcortical route to preclude any such modulation. Therefore, while our findings support a *rapid* subcortical route, they do not support a *crude* subcortical route, where “crudeness” is operationalised as specificity for low-spatial frequency information processing. On the contrary, our results provide novel causal evidence for a *generalised* functional role of the subcortical route, such that neither spatial frequency nor emotional content is filtered.

Such a mechanism is intuitive, considering that an organism’s survival is maximised if it can rapidly detect potential threats using both low *and* high spatial frequencies contained in visual stimuli^59,60^. The so-called ‘diagnostic approach’ describes flexible prioritisation of spatial frequency processing depending on the task at hand^61,62^. For example, low spatial frequency information could indicate the presence of a face but high spatial frequency information could indicate the face’s identity, either of which could be essential for detecting a potential threat^63^. The amygdala’s purported role as a ‘relevance detector’^64^ would suggest that its earliest visual input would contain all spatial frequencies and all emotional content – hence, the fast subcortical pathway unfiltered for spatial frequency or emotional content we have demonstrated in the present study. This complements other research on the auditory subcortical route to the amygdala, which was unmodulated by predictable versus unpredictable sounds^53^.

### Conclusion

Through the use of causal computational modelling, the present study has identified, for the first time, the causal functional role of the subcortical route in rapidly transmitting visual information about faces that are unfiltered for spatial frequency or emotional content. Reframing the subcortical route as playing a generalized role in rapid emotion processing unifies several opposing theories on why emotional responses to different spatial frequencies have yielded contradictory findings^6,26^; if the subcortical route transmits all spatial frequencies, as we have shown, then it may facilitate rapid neural and behavioural responses for both low *and* high spatial frequency stimuli. Thus, we propose that the supposed “coarseness” of the subcortical route may be better reframed as “unfiltered”. Future research should investigate whether the subcortical route is in fact capable of filtering spatial frequencies under certain task demands, as suggested by the diagnostic approach^61–63^, and whether this is influenced by the many cortical areas connected with the pulvinar, as suggested by previous debates on this topic^6^.

This paper makes a significant contribution to the evidence for the functional role of a subcortical route that operates in parallel with other cortical pathways for emotional visual processing. By understanding precisely what information is transmitted along this rapid, subcortical pathway and how this is used by the amygdala, we may better understand the first stages of emotional experience and the potential role subcortical activity plays in emotional disorders, such as anxiety^65^ and autism^66^.

## Methods

### Participants

Twenty-seven people participated in the study, although one was discarded due to being on psychiatric medication. This left 26 neurologically healthy participants (50% female; 23 right-handed, 3 left-handed) with an age range from 18 to 32 years (M = 22.69 years, SD = 3.87 years). All participants had normal or corrected-to-normal vision. Participation in the study was voluntary and all participants were reimbursed AUD$40 for their time. Ethical clearance was granted by The University of Queensland Institutional Human Research Ethics committee.

### Procedure

All participants were scanned at Swinburne University of Technology in Melbourne, Victoria. After removing all metal items from the body, participants were seated in the MEG inside a magnetically shielded room. Stimuli were projected onto a Perspex screen positioned approximately 1.15 meters in front of the participant (viewing angle ≈ 22.81°). Participants held a MEG-compatible 2-response button box with their dominant hand with the index and middle finger resting on the two buttons (akin to a computer mouse). The participants were instructed to fixate on the centre of the screen and remain still for the duration of each block (3 blocks of approximately 11 minutes each, with a few minutes break between blocks). A grey background was presented onscreen where faces appeared one at a time. Stimuli were presented using the Cogent 2000 toolbox for MATLAB (http://www.vislab.ucl.ac.uk/cogent.php). Each face was displayed for 200ms to minimise the likelihood of saccades. Whenever the face was not on the screen, a cross was displayed to help participants maintain central fixation. The next trial did not begin until after the participant responded using the button box to indicate whether the face was male or female. Left/right assignment for male/female did not change across the three blocks but the assignment was counterbalanced between participants. Participants were required to make their response as accurately and as quickly as they could. The inter-trial interval (ITI) was jittered between 750 and 1,500ms to reduce onset predictability.

### Stimuli

The face stimuli originated from the Karolinksa Directed Emotional Faces set (KDEF; Lundqvist, Flykt, & Ohman, 1998). Image dimensions were 198 × 252 pixels and all images were greyscale. Spatial frequency was manipulated by applying a low-pass cut-off of <6 cycles/image to create the Low spatial frequency stimuli and by applying a high-pass cut-off of >24 cycles/image to create the High spatial frequency stimuli (Fig. 1). Broad spatial frequency stimuli were images with no altered frequency information. Luminance and contrast of the Low and High spatial frequency images was matched to their respective Broad spatial frequency image by using the SHINE toolbox for MATLAB^67^. There were 60 identities: 30 males and 30 females. Each identity was presented once per condition (3 spatial frequency levels × 2 emotional expressions) resulting in six presentations per block. Hence, the three blocks resulted in a total of 180 trials per condition. All faces were presented in a random order per block but the same identity was never sequentially presented. A photodiode was placed at the bottom-left corner of the screen to record precise stimulus onset.

### MEG Data Acquisition

Neural activity was recorded using a whole-head 306-sensor (102 magnetometers and 102 pairs of orthogonally oriented planar gradiometers) Elekta Neuromag TRIUX system (Elekta Neuromag Oy, Helsinki, Finland). Activity was recorded at a sampling rate of 1,000 Hz. Before entering the MEG room, head position indicator (HPI) coils were positioned on each participant (three on the forehead and one behind each ear). An electromagnetic digitizer system was used to determine the location of the coils relative to anatomical fiducials at the nasion and at the left and right prearuicular points (FastTrak, Polhemus, Colchester, VT, USA). Electrooculographic (EOG) electrodes were also placed above and below the right eye to record eye blinks. When seated in the scanner, participants were positioned so that the helmet of the MEG was in as much contact with the head as comfortably possible. Total scanning time was 30–40 minutes, including breaks. Head position was tracked continuously using the HPI coils.

### MEG Preprocessing

The temporal extension of Signal-Space Separation (tSSS; Taulu & Simola, 2006) was applied using the MaxFilter software (Elekta Neuromag, Helsinki, Findland), actively cancelling noise and interpolating bad channels. The MaxMove function was also used to correct for head movement using a standard reference head position (0x, 0y, 40z). All subsequent offline preprocessing was completed using SPM12 (Wellcome Trust Centre for Neuroimaging, University College London, UK) via MATLAB 2014a (The Mathworks Inc., Natwick, MA, USA). The data were down-sampled to 200Hz and a band-pass filter of 0.5 – 30Hz was applied. Each participant’s MEG data were coregistered with their anatomical T1 MRI image, which was acquired at the same site (Swinburne University of Technology, Melbourne, Australia) immediately after completing the MEG scan.

Due to a technical error, EOG activity for the first four participants was not recorded. Thus, for blink correction, a frontal MEG planar gradiometer (channel MEG0922) was substituted as the EOG, given its proximity to the upper EOG sensor and the presence of the eyeblink artefact in the signal. Eyeblink artefacts were identified as a 3fT/mm deviation in the EOG signal and then marking a −300 to 300ms time window around this deviation. The associated sensor topographies were used to correct the MEG data.

For analysis, the MEG data were segmented into −100 to 650ms blocks of planar gradiometer activity time-locked to stimulus onset and baseline-corrected, creating a series of event-related fields (ERFs). For the spatiotemporal analysis, the ERF for each trial was converted to 3D scalp/time images (*x* space, *y* space, *ms* time) and then smoothed with an 8mm × 8mm × 20ms FWHM Gaussian kernel to accommodate for intersubject variability. For dynamic causal modelling, ERFs were averaged using the robust averaging function in SPM12, which weights the contribution of an epoch to the average based on its relative noise^68^. A low-pass filter of 30Hz and baseline correction were re-applied to the averaged ERFs to account for any high-frequency noise introduced by robust averaging.

### Dynamic Causal Modelling

Dynamic causal modelling (DCM) was used to compare different plausible neural networks that may underlie the observed neural data. DCM is a biologically-informed computational method of estimating the effective connectivity between brain regions using Bayesian statistics^69^. We constructed several anatomical models that were based closely on previous work by Garvert et al.^14^ who also investigated neural networks of emotional face perception. These anatomical models (Fig. 4) included the lateral geniculate nucleus (LGN; MNI coordinates: left = −22 −22 −6, right = 22 −22 −6) and pulvinar (PUL; MNI coordinates: left = −12 −25 7, right = 12 −25 7), each receiving visual sensory input, and the primary visual cortex (V1; MNI coordinates: left = −7 −85 −7, right = 7 −85 −7) and amygdala (AMY; MNI coordinates: left = −23 −5 −22, right = 23 −5 −22). All models encompassed only the first 300ms (i.e. 0–300ms post-stimulus onset) of the ERFs, as we were interested primarily in the earliest stages of visual processing.

Bayesian model estimation (via the expectation maximisation algorithm) and random-effects Bayesian model selection (which compromises between explanatory power and model complexity and accounts for variability across participants) was used to determine the most likely anatomical model for our data^70^. This was followed by a series of functional models for how these connections were modulated by either spatial frequency, emotion, or both (Fig. 5). Finally, sensitivity analysis was conducted to investigate the contribution of each parameter – either by an incremental change in an anatomical connection or by the specific modulations of spatial frequency or emotion (Fig. 6).

